# Parallelized Brightfield and Fluorescence Imaging of Organoids Using a Scalable Multi-Camera Platform

**DOI:** 10.1101/2025.10.07.681020

**Authors:** Kanghyun Kim, Rubal Singla, Amey Chaware, Jieun Park, Ina Klockner, Josh Lerner, Kevin Li, Fanghong Shen, Clay Dugo, Paul Reamey, Aurélien Bègue, Mark Harfouche, Jason L. Stein, Roarke Horstmeyer

## Abstract

Organoid viability, maturation, and growth is commonly assayed through brightfield and fluorescence microscopy using a single objective lens. However, standard microscopic imaging systems pose significant limitations for high-throughput applications, particularly in large-scale experiments where simultaneous imaging of multiple organoids requires increased throughput. There is a strong need for systems that can capture organoid growth rapidly and consistently while minimizing disturbances to culture conditions. Here, we present a novel multi-camera array scanner (MCAS) that parallelizes imaging through the simultaneous use of 48 objective lenses and sensors, resulting in a 95% reduction in acquisition times compared to commercial high-content imagers. We demonstrate and validate this system in multiple well plate formats, in both 2D and 3D neural cell cultures, and in brightfield and fluorescence. The MCAS improves efficiency for measuring organoid growth rates, assessing responses to morphogens and drugs, and measuring viral transduction efficiency. Together, these findings establish the MCAS as a scalable and versatile imaging platform for rapid phenotyping in organoid research.

## Background

Cortical organoids are three-dimensional (3D) cellular structures cultured *in vitro* that recapitulate key cyto-architectural and functional features of the developing human cortex [1]. Their value lies in their ability to capture human-specific cortical development from a given individual as well as experimental control for genetic or drug manipulations [2,3]. For example, brain organoids harboring mutations in macrocephaly or microcephaly associated genes recapitulate expected size differences as measured through brightfield microscopy, demonstrating their fidelity for modeling brain growth phenotypes [4,5]. Similarly, cortical organoid cross-sectional area is associated with inter-individual differences in the size of the infant cortex even in the absence of large effect size mutations [6]. Organoids are also useful to study teratogenic exposures, like valproic acid, that alter brain development and size [7].

Throughout the process of cortical organoid differentiation, imaging plays an important role in both quality control and phenotyping. Brightfield imaging helps to evaluate organoid survival and neuroepithelial bud structure, allowing assessment of expected differentiation outcomes, as quality control measures [6,8]. Brightfield imaging also allows evaluation of inter-individual variability or exposure-induced changes in organoid size. However, brightfield imaging alone cannot capture cell-type specific information and dynamic processes at a cellular level. Fluorescent imaging fills in the gap by enabling visualization of specific cell types or long-term tracking of clonal expansion using fluorescent reporters like GFP introduced into the system [9]. Additionally, because organoid differentiation protocols require transferring organoids between different standardized well plate formats such as 96 well plates, 90 mm dishes and 24 well plates, there is a strong need for imaging systems that can accommodate varying sizes and formats [10,11]. Organoid experiments therefore require a flexible, high-throughput imaging system capable of capturing both brightfield and fluorescence signals across organoid sizes and large dish and plate formats.

Brightfield imaging is commonly conducted using standard inverted microscopes with a single objective and sensor. While suitable for individual organoid measurements, these systems are inefficient for large-scale studies with many organoids encompassing multiple conditions, participant cell lines, or technical replicates. Although high-content plate imagers offer automated, volumetric multi-well imaging across multiple fields of view, acquisition from a full well plate can still take more than 60 minutes [12,13], and often more, as only one objective is available to axially scan at each imaging field in a step-by-step manner. A prolonged imaging time increases the duration organoids are kept outside optimal culture conditions, potentially inducing stress responses and raising contamination risk. Moreover, information regarding temporal dynamics assayed simultaneously across many wells (i.e., the observation of fluorescent reporters of calcium activity like GCaMP) is not possible in imaging systems with one objective.

To overcome these challenges, we introduce a **multi-camera array scanner (MCAS)** optimized for rapid, high-resolution imaging of thick organoids over a large field of view. The novel imaging system comprises 48 individual image sensors, each paired with a custom 0.3 numerical aperture (NA) lens providing a full-pitch resolution of 1.2 μm, comparable to a standard 10X magnification objective lens. Building on our previously published platform [14–16], we have produced an MCAS in an inverted configuration that accommodates joint capture of volumetric bright field and fluorescence images via rapid focal stack capture. This allows for parallelized acquisition of 48 full organoid volumes directly from standard culture plates. An automated stage enables lateral scanning for large organoids and movement to the second half of a 96 well plate for 96 organoid volume capture. This configuration enables volumetric imaging (multiple z-planes) of a full 96-well plate of organoids in under 26 seconds. To complement this hardware advancement, we have developed a machine learning–based segmentation pipeline built on previous work [17,18], enabling automated analysis of organoid growth, and drug response.

In this study, we demonstrate the versatility of the MCAS imaging system across multiple organoid applications. First, we monitor organoid growth and development over time, comparing various seeding conditions and media compositions to identify parameters that enhance survival. We further evaluate the impact of drugs and morphogens known to modulate progenitor proliferation and measure longitudinal organoid growth. The MCAS system accommodates any well plate format, enabling longitudinal imaging throughout all stages of organoid differentiation, highlighting its flexibility. Additionally, we showcase the system’s fluorescence imaging capabilities by quantifying varying proportions of fluorescently labeled cells in organoids and measuring fluorescence in adherent neuronal cultures. Overall, we demonstrate the efficiency and utility of the MCAS system for high throughput organoid experiments.

## Results

The MCAS comprises 48 CMOS image sensors (color, 13 Megapixel each, Onsemi AR1335) that can synchronously acquire images and video (within a 6 µs offset), enabling high-throughput, parallelized microscopy (Figure 1a). An RGB LED array provides green illumination (519 nm peak wavelength) for brightfield and blue excitation (465 nm peak wavelength) for fluorescence. Filters were used only during fluorescence imaging: an excitation filter was placed in front of the LED array and an emission filter atop the lens array. In experiments combining brightfield and fluorescence, the emission filter remained in place for both modalities because removing it required taking off the well plate; the green light could pass through the emission filter, so removal was unnecessary. Automated stages enable lateral scanning and z-stack acquisition, allowing imaging of organoids larger than a single field of view (FOV) (Figure 1b). The MCAS captures fluorescence and brightfield images across the entire 48-lens array (Figure 1c). The system supports multiple well-plate formats, including 96- and 24-well plates, providing flexibility across organoid differentiation (Figure 1d). For automated analysis, 3D scans of each organoid are converted to maximum-intensity projections to reduce dimensionality; these projections are then used for segmentation and subsequent quantitative analysis (Figure 1e).

**Figure 1.**
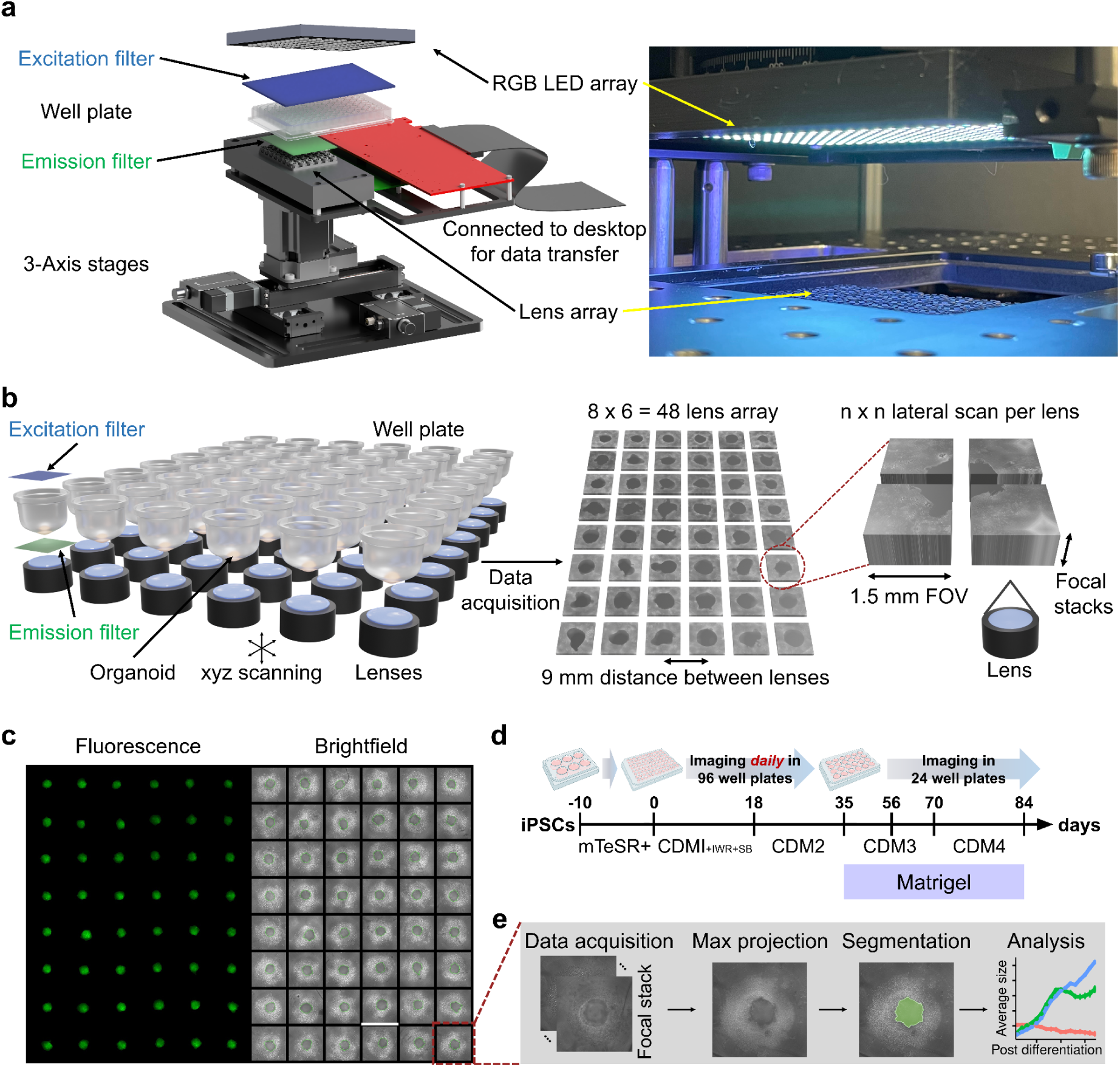
Architecture and imaging workflow of Multi-Camera Array Scanner (MCAS) for high-throughput organoid imaging. **a**. Schematics of the MCAS imaging system: The MCAS features a lens array capable of both brightfield and fluorescence imaging, together with three-axis stages that enable rapid 3D scanning of well plates. Excitation and emission filters (blue and green labeled) are utilized solely for fluorescence imaging. **b**. Each lens images an individual well, enabling a rapid, parallel 3D scan of the entire well plate. Each lens has a 1.5 mm^2^ field of view (FOV), capable of scanning a lateral grid of any given size (n × n) and capturing focal stacks at each scan location depending on the organoid size. **c**. An example image showing cortical organoids in a 96-well plate using both brightfield and fluorescence imaging. Scale bar: 1.5 mm. **d**. The MCAS system accommodates multiple plate formats, which enables longitudinal imaging throughout all stages of organoid differentiation, displayed using the protocol from [11], highlighting its flexibility. **e**. A brief workflow of imaging and analysis is as follows: 3D scans from each organoid are maximum intensity projected to reduce dimensionality. The projection is then used for segmentation, followed by quantitative analysis of cross-sectional area (y-axis) across time (x-axis) or line/exposure (color).

### Comparative analysis of MCAS and commercial systems

Each lens-sensor pair in the MCAS system covers a field of view (FOV) of 1.5 mm^2^ and a 9 mm center-to-center lens spacing, matching the well spacing of a standard 96-well plate. Lateral scanning and image stitching can be used to fully capture specimens larger than this FOV within each well [15]. The number of lateral scans varied depending on the organoid size and the well plate type. Moreover, because the MCAS contains 48 lenses, imaging a full well plate of any format requires only two scans, with each scan covering half of the plate. Due to the large, three-dimensional structure of cortical organoids, axial scanning was performed across all experiments. Thick tissues like organoids exhibit substantial variation in focal depth across different regions, and a single focal plane often fails to capture their true boundaries. To avoid inaccurate estimation of organoid size and loss of structural information, multiple focal planes are therefore acquired and later combined, ensuring that sharp outlines are retained across varying depths [19]. 3D data acquisition time varies depending on the number of lateral scans and the number of focal planes used. Figure 2b compares acquisition times across use cases under brightfield illumination. The MCAS achieves rapid throughput in brightfield by using a continuous, on-the-fly axial scanning strategy: images are captured and written to disk while the Z-stage moves at constant velocity, eliminating the stop-and-settle overhead of conventional step-and-shoot approaches. In contrast, the same on-the-fly strategy is not applicable to fluorescence imaging. Fluorescence typically requires longer exposure times to achieve adequate signal-to-noise ratio, so continuous motion during fluorescence z-stack capture can introduce motion blur. To maintain spatial fidelity, fluorescence stacks were therefore acquired in a step-and-settle mode: the stage moves to each axial plane, settles, the sensor integrates for the prescribed exposure, and then advances to the next plane. As a result, while brightfield volumes can be scanned quickly with continuous motion, volumetric fluorescence acquisitions require additional time due to image exposure and step-and-settle constraints.

**Figure 2.**
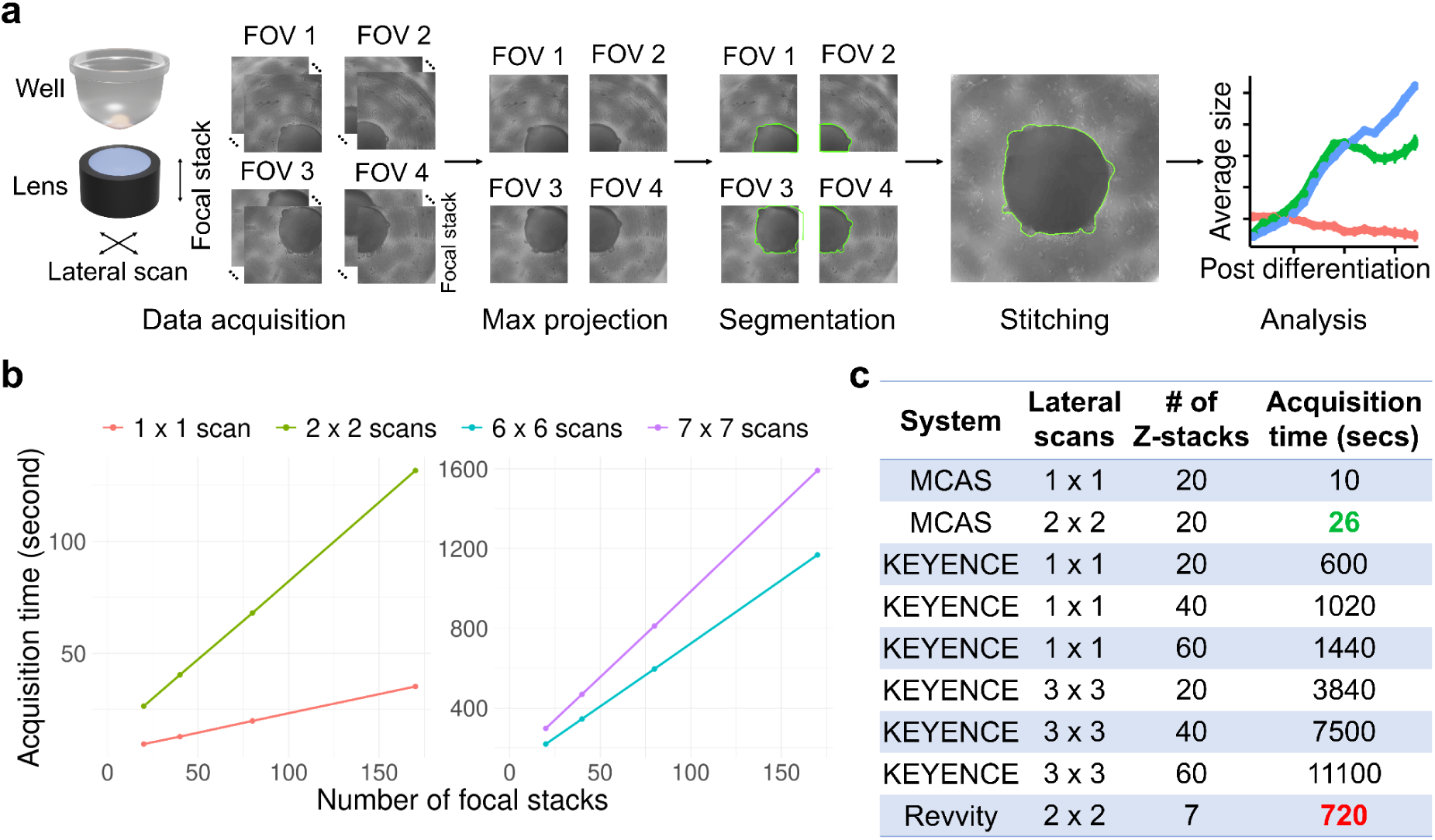
Overview of the MCAS pipeline and a data acquisition time comparison. **a**. A lateral and axial scan is performed to fully image organoids in 3D. Each field of view (FOV) is maximum projected and segmented individually. The segmented organoid images are then stitched together, and the stitched images are used for subsequent analysis. Green outlines indicate the segmentation boundaries of organoids. **b**. The colored lines show the 3D data acquisition time for a 96-well plate as a function of the number of lateral scans and focal stacks using the MCAS, based on brightfield imaging. The four reported lateral scan settings are the ones we used in our experiments. A 7 × 7 lateral scan is performed to fill FOV gaps between lenses, enabling full well-plate imaging without gaps between wells when necessary. **c**. Acquisition time comparison between the MCAS, the KEYENCE BZ-X710, and the Revvity Opera Phenix® Plus, demonstrating a 95% reduction in acquisition time (from 720 seconds (12 minutes) to 26 seconds).

To estimate the acquisition time for imaging a 96-well plate in brightfield relative to commercially available whole-plate scanning microscopes (high-content imagers), we experimentally measured acquisition times on two commercial systems. Figure 2c compares acquisition time across MCAS, the KEYENCE BZ-X710, and the Revvity Opera Phenix® Plus as a function of the number of lateral scans and focal (Z) stacks. MCAS completed a scan in 26 seconds, representing a 95% reduction in acquisition time compared with 720 seconds (12 minutes) on the fastest available commercial system (Revvity). The KEYENCE BZ-X710 took at least 600 seconds for focal stack acquisition alone, and the time can extend to several hours when lateral scanning is also required.

### Using MCAS to determine the impacts of differentiation optimizations

Cortical organoid differentiation involves seeding a specific number of pluripotent cells in non-adherent dishes, followed by the use of different media, signaling molecules, and growth factors to guide the differentiation toward expected cell types. We tested modifications of a reproducible cortical organoid differentiation protocol [11] to enable consistent differentiations with limited cell death across iPSCs from multiple participants (Figure 3a). The original organoid differentiation protocol describes seeding cells in cortical differentiation media 1 (CDM1) media at Day 0 [11]. However, we found some iPSC lines had limited growth when transitioned directly from pluripotency media (mTeSR Plus, StemCell Technologies) at the iPSC stage to CDM1 media for seeding. To find the seeding condition that leads to high survival and growth for multiple iPSC lines (details in Methods section), we tested three conditions in an iPSC line that failed to differentiate well with the published protocol [11] and evaluated their impact on organoid viability and growth using the MCAS: seeding directly in CDM1 as per the original protocol (Condition 1), seeding in mTeSR Plus followed by half of media change with CDM1 on Day 1 (Condition 2), and seeding in mTeSR Plus followed by replacement of three-quarters of the media with CDM1 on Day 1 to provide a gradual transition (Condition 3).

**Figure 3.**
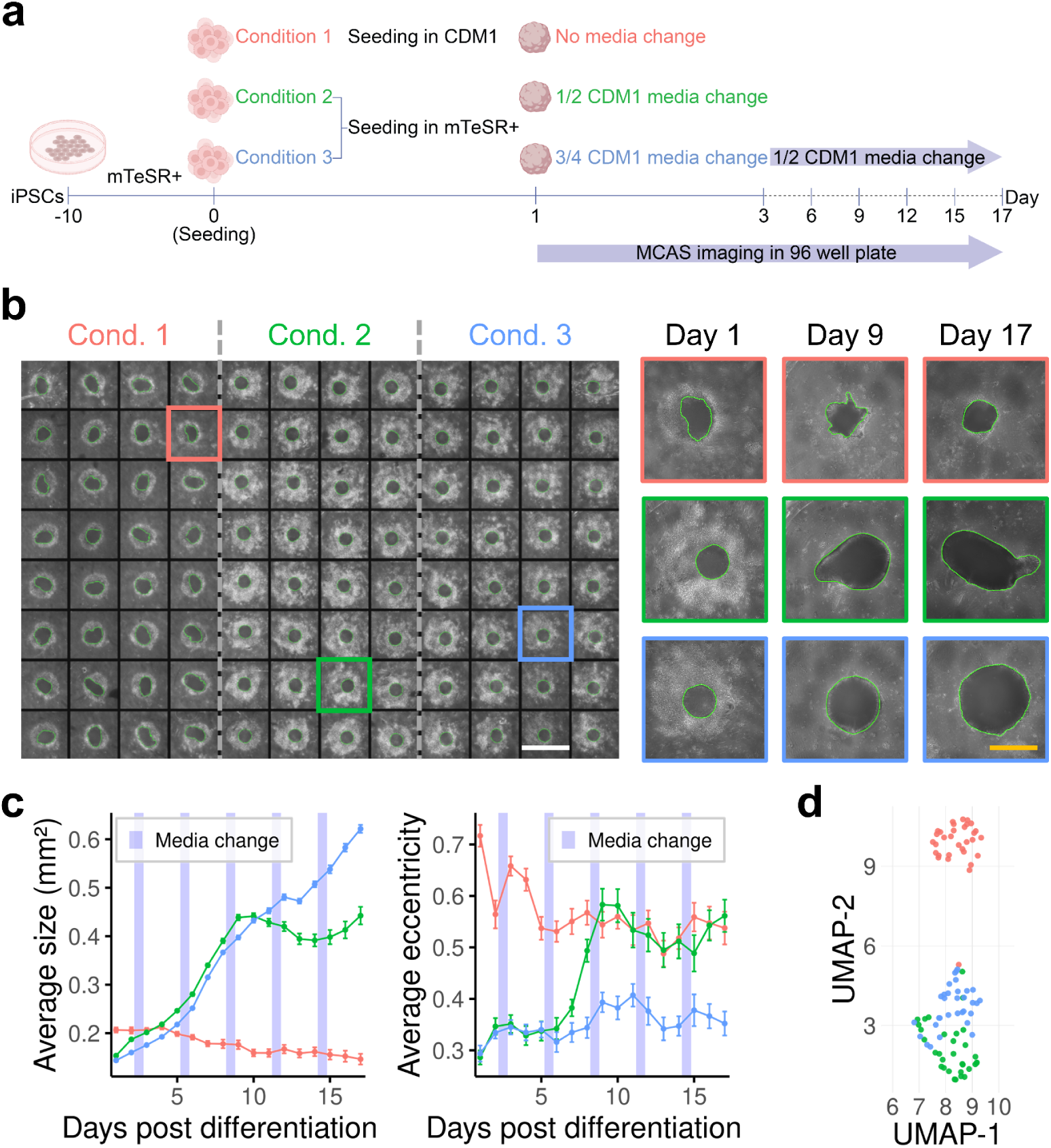
The impact of seeding conditions on longitudinal organoid growth assayed through the MCAS. **a**. To determine the optimal seeding condition for maximizing survival across iPSC lines, three conditions were tested in an iPSC line that exhibited poor differentiation with the published protocol [11] assessing their effect on organoid viability and growth. **b**. Example images for each condition taken over 17-days. Green outlines indicate the segmentation boundaries of organoids. Left scale bar: 1.5 mm. Right scale bar: 0.6 mm. **c**. Longitudinal cross-sectional area and eccentricity highlight the differences in organoid growth patterns across conditions. Error bars indicate standard error from the mean. 30, 31, and 31 organoids were imaged for condition 1, 2, and 3, respectively. **d.** UMAP plot of CLIP embeddings on Day 9, showing clear condition-specific clusters driven by image texture differences. Each dot represents an organoid, colored by condition.

Organoids were imaged daily in a 96-well plate from the day after seeding (Day 1) through Day 17. We observed that organoids seeded in CDM1 alone (Condition 1) decreased in cross-sectional area between Day 1 and Day 17 (Figure 3c). However, organoids seeded in mTeSR Plus media (Conditions 2 and 3) exhibited higher growth compared to those seeded in CDM1 alone (Condition 1) (Figure 3c). Condition 3 led to consistent growth over time, indicating that the timing and initial media composition play a crucial role in organoid differentiation outcomes.

In addition, the cross-sectional images of organoids in Condition 3 were more circular, or equivalently had lower eccentricity, which is generally a feature of on target differentiation. In contrast, organoids in Condition 1 with stalled growth showed high eccentricity (Figure 3c). A similar trend was also observed in Condition 2, where organoids began displaying an irregular growth pattern on Day 9. Notably, eccentricity in Condition 2 started increasing on Day 7, two days prior to the stall in growth suggesting that eccentricity may serve as an early indicator of unexpected organoid differentiation (Figure 3c). To investigate potential textural or morphological differences across conditions, we obtained CLIP [20] embeddings of the MCAS-obtained images of the organoids on Day 9 and visualized them using a 2D UMAP [21] scatter plot. The plot revealed clear condition-specific clusters (Figure 3d), providing further evidence for differences in organoid structure across conditions. After Day 14 of differentiation, eccentricity values became more variable across all conditions. A likely explanation is that organoids begin to develop neuroepithelial buds, mimicking early neural tube structure, around Day 14 resulting in less circular shapes. Therefore, eccentricity could serve as a reliable predictor of organoid growth, but only up to Day 14. Overall, we see that lower eccentricity corresponded to more circular organoids and was associated with greater cross-sectional area. Thus, MCAS enables rapid (35 seconds to image organoids in a 96-well plate with 170 focal planes), quantitative comparison of growth and eccentricity across conditions with minimal manual intervention, unlike conventional microscopes such as EVOS.

### Imaging the impact of small molecules on organoid growth

We next tested the effects of valproic acid (VPA), a well known medication used to treat epilepsy and bipolar disorder where prenatal exposure increases risk for autism spectrum disorders and intellectual disability [22–24], CHIR99021 (CHIR), a known GSK3β inhibitor that activates the canonical Wnt signaling pathway and promotes progenitor proliferation [25–27], and Mitomycin C, an anticancer chemotherapy drug, that strongly inhibits cell proliferation [28,29]. To evaluate its effects on progenitor proliferation within cortical organoids, we tested a range of concentrations and exposure durations (Figure 4a). In our previous study, we observed opposite effects of VPA and CHIR on 2D human neural progenitor cell proliferation: VPA decreased while CHIR increased proliferation [30]. To demonstrate the utility of the MCAS system for rapid longitudinal exposure screening, we imaged over two weeks of differentiation the effect of different concentrations of these small molecules with known impacts on proliferation.

**Figure 4.**
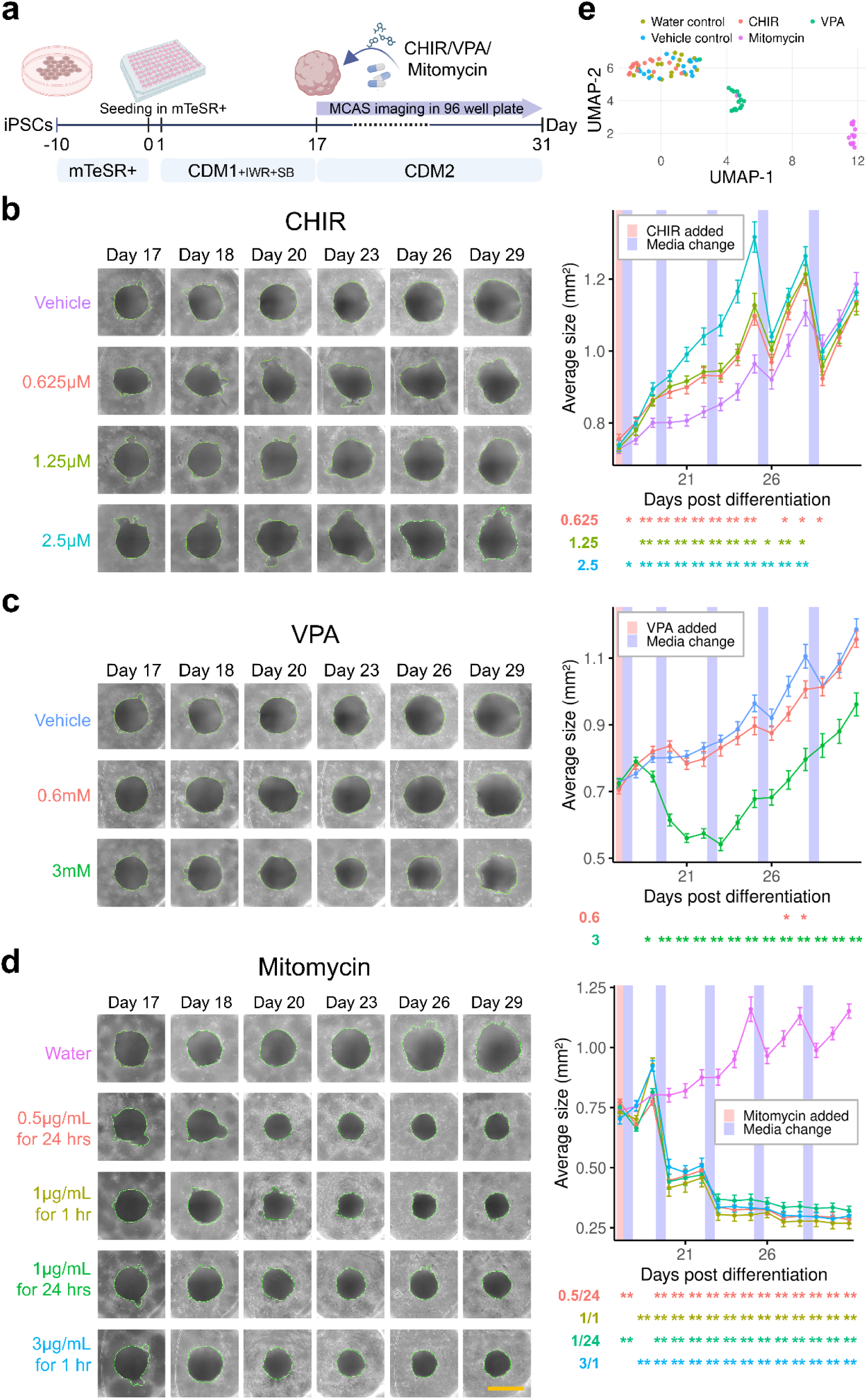
The impact of small molecules on cortical organoid growth. **a**. We tested the effects of multiple concentrations of valproic acid (VPA), CHIR99021 (CHIR), and Mitomycin C on organoid cross-sectional areas from Day 17 to Day 31. **b-d**. Cross-sectional area curves for each small molecule and concentration are reported, along with example images. Asterisks below each line plot indicate p-values from Welch’s *t*-tests comparing each drug with the control: * for p < 0.05 and ** for p < 0.01. **b** and **c** share the same vehicle control and the vehicle representative images are repeated for visualization. Green outlines indicate the segmentation boundaries of organoids. 16 organoids were imaged per condition. Error bars indicate standard error from the mean. Scale bar: 0.93 mm. **e.** UMAP plot of CLIP embeddings on Day 22, showing clear condition-specific clusters. The clusters, from left to right, correspond to CHIR (2.5 µM), water and vehicle controls, VPA (3 mM), and Mitomycin (0.5 µg/mL for 24 hours) treatments, respectively. The colors in **e** are not the same as those in **b**-**d**.

The graphs in Figure 4b-d highlights the effects of CHIR, VPA, and Mitomycin C exposures on the cross-sectional area of organoids compared to the vehicle control group from Day 17 to Day 31. All CHIR-treated groups (0.625 µM, 1.25 µM, and 2.5 µM) exhibited a dose-dependent increase in organoid cross-sectional area (Figure 4b). The 2.5 µM CHIR group showed the largest increase, with a 13% larger area compared to the vehicle control on Day 26. Similarly, the 1.25 µM and 0.625 µM CHIR groups resulted in approximately 9% and 5% larger areas, respectively, compared to the control. Notably, transient decrease in organoid cross sectional area was observed immediately following each media change (imaged just after media change) for all conditions. This reduction likely reflects the removal of cellular debris surrounding the organoids during media change rather than a true biological reduction in organoid size.

In contrast, VPA-treated groups experienced a significant reduction in cross-sectional area. The 3 mM VPA group showed the greatest decrease, with areas 26% smaller than the control on Day 26 (Figure 4c). The 0.6 mM VPA group had a moderate reduction in size, maintaining areas 5% smaller than the vehicle control. Mitomycin-treated groups exhibited a significant decline in organoid growth, which varied with concentration and exposure duration. Specifically, Mitomycin at 1 µg/mL for 24 hours and 3 µg/mL for 1 hour resulted in a decrease of approximately 37-45% in organoid size relative to the control by Day 20, with no recovery observed thereafter (Figure 4d). Treatment with 0.5 µg/mL for 24 hours showed a milder but consistent reduction, with organoid sizes approximately 44% smaller than the control. The results reveal distinct effects of Mitomycin on organoid growth, emphasizing the influence of treatment. Although Mitomycin led to a significant decrease in organoid growth compared to the control, we did not observe a clear time-sensitive or dose-dependent effect.

As before, to further investigate the morphological or textural differences, we generated CLIP embeddings of the MCAS-obtained images on Day 22, and visualized them using a 2D UMAP scatter plot (Figure 4e). Organoids treated with VPA (3 mM) and Mitomycin (0.5 µg/mL for 24 hours) formed distinct clusters. Although CHIR-treated organoids (2.5 µM) were near the same overarching cluster as the controls, they were mostly separable from the vehicle and water control groups. As expected, the vehicle and water controls were intermingled. We also found clear, condition-specific clusters, further demonstrating the impact of these small molecules on organoid structure.

Overall, the MCAS allowed rapid imaging, even while performing 2 × 2 lateral scanning to image the full size of the organoids (132 seconds to image organoids in a 96-well plate with 170 focal planes). This demonstrates the ability of MCAS to rapidly phenotype organoid size and structure in response to small molecules, showing expected and concentration-dependent changes over time.

### Rapid imaging of fluorescence in organoids

To evaluate the utility of the MCAS system for measuring fluorescently labeled organoids, we we generated cortical organoids by mixing iPSCs transduced with lentivirus expressing a constitutively active eGFP reporter [31] with non-transduced iPSCs at varying ratios (20%, 40%, and ∼100%) (Figure 5a). Organoids were imaged over a seven-day period using the MCAS imaging system.

**Figure 5.**
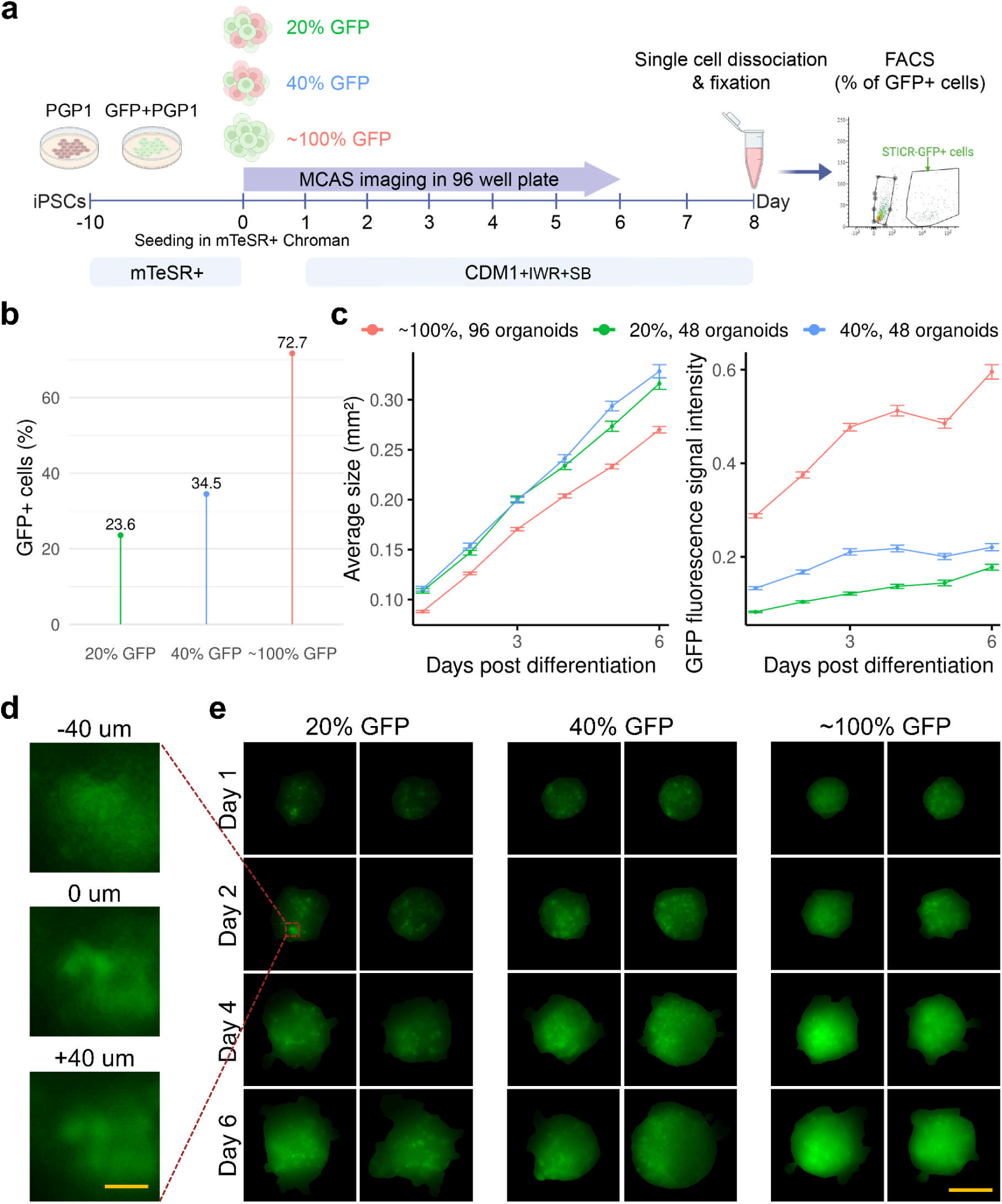
Rapid fluorescence imaging of organoids via the MCAS. **a**. We generated cortical organoids by mixing iPSCs transduced with a constitutively active GFP reporter with non-transduced iPSCs at varying ratios (20%, 40%, and ∼100%). **b**. FACS sorting results for each condition. **c**. We imaged the organoids from Day 1 to Day 6 and show the cross-sectional area growth and the fluorescence signal intensity over time using MCAS imaging. Higher transduction limits organoid growth but results in increased fluorescence. **d**. Thick organoids necessitate acquisition of multiple focal planes followed by maximum intensity projection for fluorescence signal intensity analysis. Scale bar: 0.33 μm. **e**. Example images for each condition and differentiation times are shown. Scale bar: 350 μm.

Brightfield imaging showed a gradual increase in the cross-sectional area of organoids across all three conditions during the 6-day differentiation timeline, suggesting that lentivirus exposure does not halt organoid growth. However, organoids in the ∼100% condition consistently exhibited smaller cross-sectional areas compared to the 20% and 40% conditions (Figure 5c (left)). By Day 6, the organoid area in the ∼100% condition was approximately 17-22% smaller than in the 20% and 40% conditions, which were comparable in size. This observation indicates that a higher proportion of lentivirus (∼100% condition) may lead to reduced organoid growth. The smaller size of organoids in the ∼100% condition aligns with previous reports that lentiviral vectors can negatively affect proliferation [32].

Figure 5c (right) shows the fluorescence signal intensity, reflecting the efficiency of eGFP transduction. The 20% condition displayed the lowest fluorescence signal throughout, increasing steadily from Day 1 to Day 6 with minimal variability. As expected, the fluorescence intensity values in this condition were consistently lower than in the other conditions due to the lower proportion of transduced cells. The 40% condition showed fluorescence signal intensities approximately twice as high as those in the 20% condition, with a similar trend of steady increase and slightly higher variability. The ∼100% condition exhibited the highest fluorescence signal intensity, with values approximately 3-4 times higher than the 20% condition and twice higher than the 40% condition on average. These findings confirm a proportional relationship between the percentage of transduced cells and fluorescence signal, with the ∼100% condition demonstrating the strongest signal due to the maximum number of transduced cells.

Figure 5d shows the representative z-stack images of organoids at different focal planes (−40 µm, 0 µm and +40 µm) at the second day of differentiation. The z-stack is necessary because the organoids are 3D structures with significant thickness, where a single focal plane will be blurred by GFP signal from other planes. The MCAS system acquires z-stacks which helps to reconstruct a more accurate representation of fluorescence throughout the organoid volume, reducing the risk of underestimating the transduction efficiency or spatial expression. The MCAS was able to capture the fluorescent signal, but we could not resolve individual cells, especially for ∼100% GFP, due to light scattering in dense tissue.

We further performed Fluorescence-Activated Cell Sorting (FACS) to validate the MCAS fluorescence measurements. Supplementary Figure 2 shows the gating strategy and the percentage of GFP-positive cells in each condition after sorting. FACS analysis confirmed that the proportion of GFP-positive cells closely reflected both the initial mixing ratios of transduced and non-transduced iPSCs and the fluorescence readouts obtained with MCAS. In the 20% condition, approximately one-fifth of the cells were GFP-positive. The 40% condition displayed nearly double that fraction, while the ∼100% condition showed almost complete GFP positivity, with only a small proportion of GFP-negative cells that could reflect either incomplete transduction at the iPSC stage or transgene promoter silencing during differentiation (Figure 5b). These results are consistent with the fluorescence intensity curves from MCAS (Figure 5c), in which increasing GFP-positive cell proportions corresponded to progressively stronger organoid fluorescence signals. Importantly, the concordance between FACS and MCAS highlights the utility of MCAS as a rapid, plate-based imaging platform that can serve as a reliable proxy for more labor-intensive single-cell assays, enabling high-throughput quantification of transduction efficiency in 3D organoid cultures.

The MCAS system rapidly measures fluorescence intensity under varying viral conditions, demonstrating its potential as a valuable tool for assessing multiplicity of infection (MOI) and optimizing viral delivery protocols in organoids. The time required to image organoids in a 96-well plate with 80 focal planes was 20 seconds for brightfield and 110 seconds for fluorescence. As mentioned above, fluorescence stacks were acquired in step-and-settle mode, which resulted in a longer acquisition time than brightfield.

### Quantifying transfection efficiency and confluency in monolayer cells

To evaluate the broad utility of the MCAS imaging platform for quantifying both transfection efficiency and cellular confluency in monolayer cells as well as 3D organoids, we transfected adherent Neuro2a (N2a) neuroblastoma cells with a GFP-expressing plasmid in a 96-well plate format.

Brightfield and fluorescence images rapidly acquired on the MCAS allow the quantification and colocalization of individual cells with GFP fluorescence (Figure 6a-c). Analysis of MCAS images allows quantification of variability in GFP transfection efficiency (Figure 6b) and confluency (Figure 6c) across the 96-well plate. Transfection rates, quantified by the proportion of fluorescent cells, ranged from 26.3% to 43.3% across wells. The confluency measurement was obtained via brightfield imaging and revealed a range from 11.2% to 28.9%. Together, these results demonstrate the utility of MCAS imaging for simultaneous quantification of fluorescence-based transfection efficiency and brightfield-derived confluency metrics in a 96-well plate format. This supports the feasibility of using MCAS for automated screening applications involving fluorescent reporter assays in 2D cell cultures with rapid acquisition times. The time required to image cells in a 96-well plate with 80 focal planes was 345 seconds for brightfield and 51 minutes for fluorescence. As mentioned above, fluorescence stacks were acquired in step-and-settle mode, which resulted in a longer acquisition time than brightfield.

**Figure 6.**
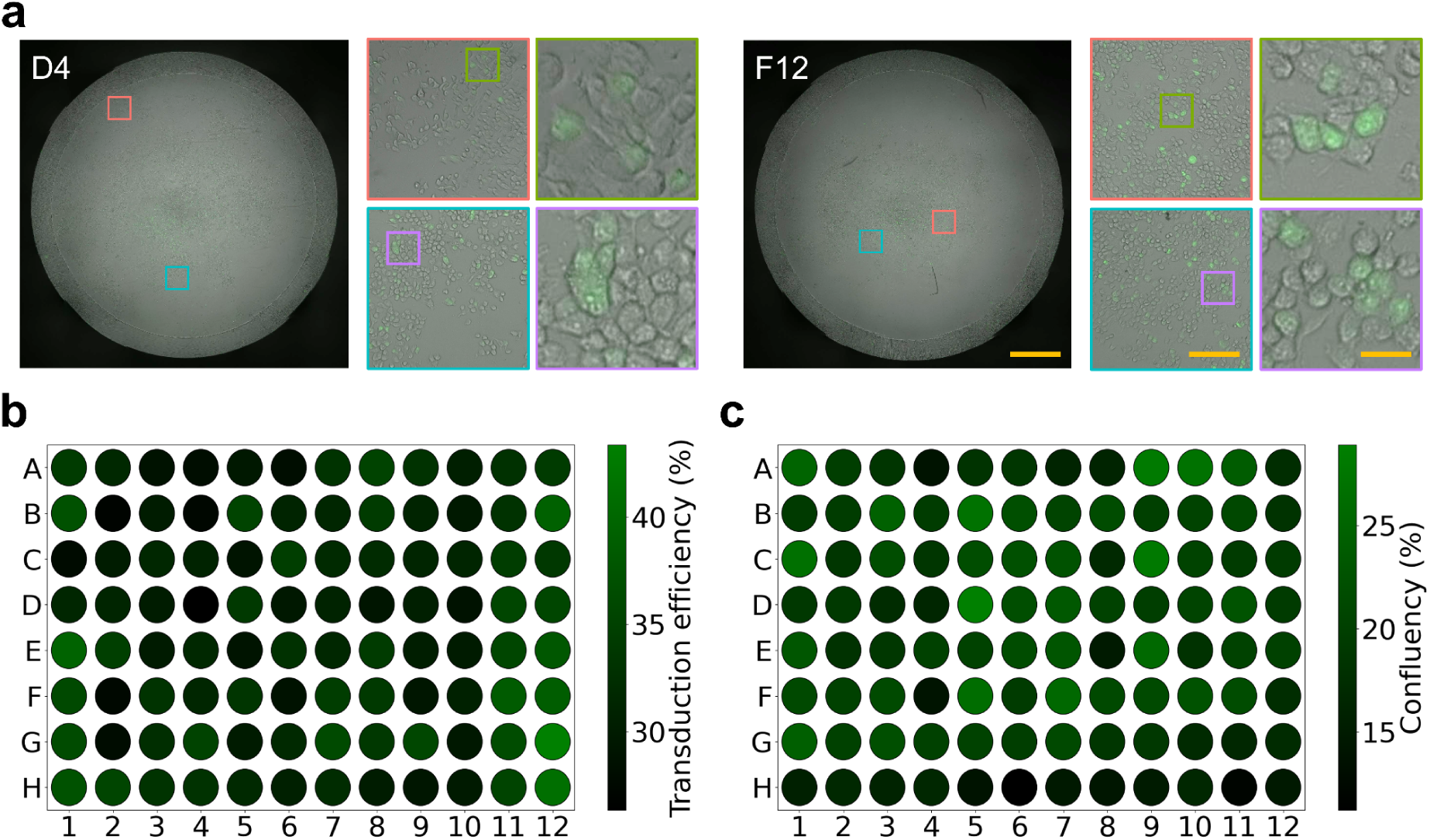
Rapid fluorescence imaging of monolayer cells via the MCAS. **a**. Example images of mouse neuroblastoma cells (Neuro2a; ATCC CCL-131) transfected with Addgene plasmid #24526, which expresses eGFP under the human ubiquitin C (UbC) promoter. Images from two separate wells of a 96-well plate are shown as zoomed images. Brightfield and fluorescence images are superimposed. Left scale bar: 1 mm. Center scale bar: 160 μm. Right scale bar: 32 μm. **b**. Transduction efficiency calculated as the number of GFP+ cells divided by the total number of cells across wells can be calculated using MCAS. **c**. Cellular confluency calculated as the total area of the wells occupied by the cells can also be assayed.

### MCAS imaging of larger, later-stage organoids in a 24-well plate format

Organoids are commonly differentiated beyond Day 50 to generate later born cells and enhance maturation. As organoids grow in size, they can no longer be cultured in 96-well plates, as the limited space may restrict their development. Organoids are transferred from 96-well plates to 90 mm dishes at Day 18, and subsequently to spinner flasks at Day 35 to support continued growth and maturation, based on a commonly used protocol [11].

Since organoids are not maintained exclusively in 96-well plates throughout differentiation, it is essential to ensure MCAS can be imaged in alternative plate formats. With conventional systems such as EVOS, imaging and quantifying later-stage organoids is challenging: individual organoids often exceed a single field of view, there is no stitching function to generate composite images, and it becomes difficult to track which organoids have or have not been imaged across multiple, unstitched frames. To address this, we evaluated the feasibility of imaging organoids at various stages of differentiation (Days 35, 56, 70, and 84) using a 24-well plate format (Figure 7a). Each well of the 24-well plate contained multiple organoids, enabling continued high-throughput imaging.

**Figure 7.**
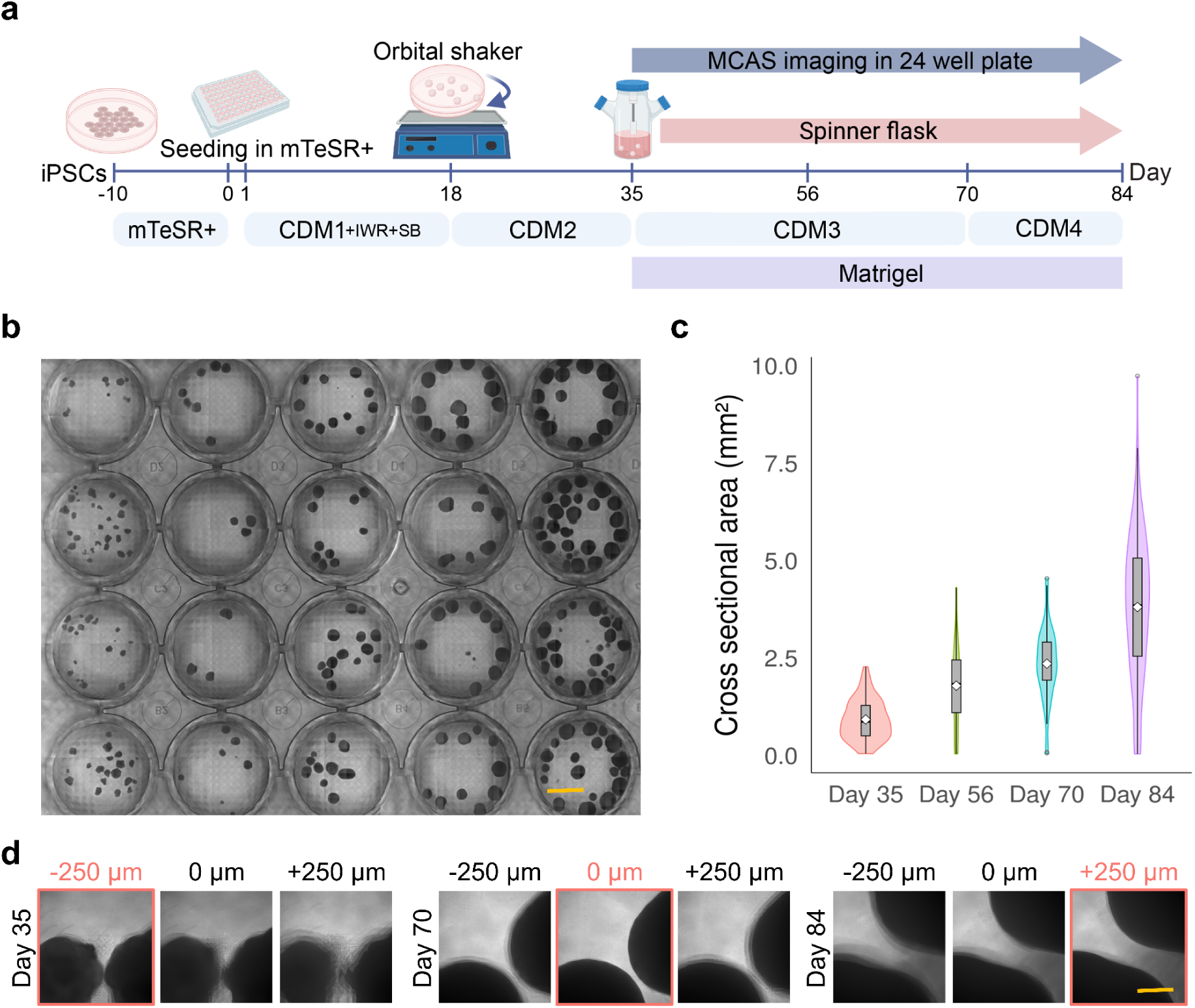
Imaging later-stage organoids in a 24-well plate. **a**. Timeline of the cortical organoids from iPSC stage, organoid differentiation starting from seeding (Day 0) to Day 84. Organoids from different iPSCs lines generated at various stages of differentiation (Days 35, 56, 70, and 84) were placed in a 24-well plate. **b.** Representative brightfield image of multiple organoids in a 24-well plate. Scale bar: 5.9 mm. **c**. Violin plots show the organoid cross-sectional area measurements per group. 77, 20, 45, and 123 organoids were imaged at Day 35, Day 56, Day 70, and Day 84, respectively. **d**. Organoids at different stages of differentiation have different sizes, requiring different axial scanning ranges. Images show the sharpest outline of each organoid in each differentiation group at different focal planes. Scale bar: 0.5 mm.

Figure 7c shows cross-sectional area measurements of organoids derived from multiple iPSC lines at multiple time points in differentiation. These organoids were generated from different iPSC lines. The organoid cross sectional area progressively increases with differentiation time. These results demonstrate that the MCAS imaging platform is adaptable with imaging organoids across multiple stages of differentiation and plate formats, providing a flexible platform for longitudinal imaging of organoid growth (468 seconds to image organoids in a 96-well plate with 40 focal planes).

Figure 7d illustrates the importance of axial (Z-axis) scanning for accurately imaging organoids at later stages of differentiation. Because organoids are large 3D structures that increase in size and thickness over time, a single focal plane cannot reliably capture their boundaries. The panel shows representative images of organoids at Days 35, 70, and 84 acquired at three different focal depths (−250 µm, 0 µm, and +250 µm). At each stage, the organoid outline is sharpest at a different focal plane, reflecting differences in organoid size. This demonstrates that later-stage organoids require broader axial scanning ranges to capture their full cross-sectional area accurately. Without Z-scanning, measurements could underestimate organoid size or miss structural features, particularly in thicker, more mature organoids.

### Data visualization

The massive volume of data produced by the MCAS necessitates the use of automated analysis tools, though occasional manual image inspection is still required. To facilitate smooth exploration of high-resolution images, we created Gigaviewer—a custom, web-based viewer built on OpenSeadragon [33]. Beyond the standard zoom and pan functionalities provided by OpenSeadragon, Gigaviewer includes user-adjustable focus controls. It generates images at multiple zoom levels to support dynamic resolution display, while also preloading neighboring fields of view to ensure fluid and continuous navigation. Example images can be found at https://gigazoom.rc.duke.edu.

## Discussion

High-content imaging is essential for both quality control during organoid differentiation and for providing inexpensive, scalable phenotyping assays. Brightfield imaging allows researchers to quickly assess organoid survival, circularity, and neuroepithelial bud formation, while fluorescence enables tracking of specific cell populations or reporter constructs. However, current imaging approaches are often too slow, require manual imaging of individual fields, or cannot accommodate organoids across different culture formats, making them poorly suited for large-scale studies.

The Multi-Camera Array Scanner (MCAS) directly addresses these limitations by enabling rapid, parallelized imaging across multiple wells and z-planes. Compared to EVOS or high-content imagers, which typically require 12-64 minutes to image a 96-well plate, MCAS completes the same task in 26 seconds, representing a 95% reduction in acquisition time. This efficiency gain was consistently demonstrated across diverse applications: (i) daily monitoring of differentiation and seeding conditions, (ii) quantifying drug-induced changes in growth, (iii) fluorescence-based measurements of viral transduction in organoids, (iv) assessing transfection efficiency and confluency in monolayer cultures, and (v) longitudinal imaging of later-stage organoids in larger plate formats. In each case, MCAS enabled both rapid acquisition and automated analysis, increasing throughput for large-scale organoid growth experiments. Moreover, unlike many commercial high-content imagers that require a dedicated space or room, the MCAS fits on a desk alongside a computer and imaging system (50 cm × 50 cm × 50 cm) and eliminates the need for well-plate transportation. Together, these results establish MCAS as a time-efficient and versatile imaging solution relative to conventional microscopes.

MCAS also enabled several biological insights across experimental contexts. Because our project required a single standardized seeding protocol across all iPSC lines, we tested multiple conditions to identify one that worked broadly. Seeding experiments demonstrated that the initial choice of media and the timing of transitions significantly influence organoid survival and growth. Using mTeSR+ media during early stages improved survival and initial expansion, likely by buffering stress from rapid media changes. The gradual transition to CDM1, as implemented in Condition 3, yielded more consistent outcomes across iPSC lines. These minor modifications of the differentiation protocol may make them more broadly applicable. Moreover, eccentricity emerged as a potential early marker of failed differentiation (Figure 3).

Drug perturbation assays further validated the system’s biological utility, recapitulating expected responses: CHIR enhanced organoid growth, VPA suppressed it, and Mitomycin induced severe growth arrest (Figure 4). These results confirm that MCAS can support rapid, high-throughput drug screening with quantitative readouts. Fluorescence-based assays demonstrated a proportional relationship between transduced cells and reporter fluorescence intensity, but also revealed that excessive viral exposure can compromise organoid development, underscoring the need to optimize viral delivery protocols (Figure 5). Finally, imaging later-stage organoids (Days 35-84) emphasized the necessity of z-stack scanning: single focal planes underestimated organoid size and missed structural features, whereas volumetric imaging captured the full dimensions of large, thick organoids (Figure 7). Collectively, these findings demonstrate that MCAS not only accelerates image acquisition but also provides a robust platform for detailed biological phenotyping of organoid development and treatment responses.

To further increase the data acquisition speed of the MCAS, we can consider three possible approaches. First, we can reduce the number of focal stacks by either increasing the axial step size or decreasing the axial scanning range. For the experiments in this manuscript, we used axial step sizes of 5-40 µm, depending on the specific task and samples. The step size could be increased further as long as it does not impact the accuracy of the cross-sectional area measurements. We also used a scanning range of 200 µm to 1.7 mm, which can be decreased to reduce the number of focal stacks. Developing a method to automatically determine the optimal step size and scanning range without sacrificing measurement accuracy is an area for future work. Second, we can increase the acquisition speed by using stronger fluorescence excitation sources. Longer exposure times were the main bottleneck in acquisition speed. Using a more powerful LED or a laser to increase excitation light intensity could significantly improve imaging speed, which is essential for functional assays such as calcium imaging with reporters like GCaMP, that we plan to explore in future work. Finally, we can consider using a monochrome sensor instead of a color (Bayer) sensor to improve quantum efficiency for fluorescence measurements, thereby reducing exposure time and overall scan time.

Although MCAS offers substantial speed gains, several limitations remain. Lateral scanning could not capture the full extent of a 24-well plate, leading to cropped organoids in the rightmost column in a 24-well plate. This limitation could be mitigated by either performing separate scans of missing regions, at the cost of additional time, or by demagnifying each lens to expand the field of view—a modification that would require minor adjustments to the MCAS lens array layout. Another issue we encountered during 24-well plate imaging was stitching artifacts. These artifacts were mainly due to two reasons. First, the 24-well plate has a larger surface area per well compared to 96-well or higher-density plates, which made the organoids more susceptible to shifting from stage movement vibrations. This introduced lateral shifts in the stitched images. Second, we placed later-stage organoids in the 24-well plate, which appeared dark during imaging due to their increased thickness. The stitching algorithm struggled in these cases due to a lack of distinctive features needed for accurate alignment. Finally, the system currently supports only GFP fluorescence and lacks additional light sources for multichannel fluorescence imaging.

Hardware and software improvements could further enhance MCAS performance. Widefield illumination for fluorescence imaging leads to reduced contrast due to out-of-focus signal; this could be mitigated by integrating light-sheet microscopy [34,35] which would also reduce phototoxicity, though at the expense of throughput since some lenses would need to be replaced with illumination optics. Another potential hardware improvement is the integration of stage-top incubation systems or enclosed incubators into the MCAS, which would help maintain optimal culture conditions during extended imaging sessions. Lastly, although image acquisition is rapid, data processing and analysis were performed post-acquisition. We believe that real-time analysis is achievable by processing data concurrently with acquisition. In other words, while the MCAS scans one location, data from the previous location can be analyzed. We plan to demonstrate this capability in future work.

### Conclusion

As organoid studies expand in scale and complexity, ranging from patient-derived iPSCs to high-throughput drug and genetic screens, the demand for **fast, flexible, and quantitative imaging systems** will continue to grow. The MCAS platform addresses this demand by combining speed, scalability, and adaptability to multiple plate formats.

## Methods

### MCAS methods

Here, we discuss the workflow for MCAS data acquisition, image processing, and analysis, which is commonly used across experiments. Each experiment utilized slightly different settings tailored to its specific imaging and processing requirements. Detailed settings for each experiment are provided in the corresponding method sections.

### MCAS hardware

For illumination, we used an 8 × 12 LED array with a 9 mm pitch (96-well RGB LED microplate light panel from Tindie), positioned approximately 6.5 cm above the sample plane. The LED array consists of RGB LEDs (SK6812) with separate red, green, and blue emitters. The peak wavelengths are 629 nm, 519 nm, and 465 nm for the red, green, and blue LEDs, respectively. The wavelength spectra of these LEDs are shown in [36]. The blue channel is used to excite GFP, so during fluorescence imaging only the blue channel is activated. For brightfield imaging, only the green channel is used. A circular baffle was placed beneath the LED array to ensure each specimen was illuminated solely by the LED directly above it, enhancing image contrast. To scan specimens laterally and axially, we employed three motorized linear stages from Zaber (X-LSM-E Series with travel distances of 100 and 50 mm, and X-VSR-E Series with a travel distance of 20 mm). Details on the primary optical characteristics of the customized lenses used in our MCAS can be found in our previous publication [14]. For fluorescence imaging, an ET470/40x excitation filter (80 × 100 mm²) and an ET525/30m emission filter (52 × 70 mm²) from Chroma Technology were used. Prior to fluorescence imaging, the excitation filter was placed over the plate, and the emission filter was positioned on top of the lens array. When imaging GFP-expressing specimens in brightfield, only the excitation filter needs to be removed. This is much easier than removing the emission filter, which is mounted directly above the objectives. The LEDs’ green channel passes through the emission filter because its spectrum overlaps the GFP emission band; therefore, removing the emission filter is unnecessary.

### MCAS data acquisition

The MCAS uses motorized stages to perform lateral and axial scanning. When performing lateral scans, we used a step size of 1.3 mm to create overlaps between adjacent FOVs for seamless image stitching. For axial scanning, a range of 200 µm to 1.7 mm was employed, with step sizes of 5, 10, or 40 µm. In cases where additional GFP signals are needed, we perform a separate imaging session using appropriate excitation and emission filters, followed by brightfield imaging. When imaging and media changes occurred on the same day, imaging was always performed after the media change.

### Comparison of MCAS and Commercial Systems

We experimentally compared the acquisition time between the MCAS, the KEYENCE BZ-X710, and the Revvity Opera Phenix® Plus. An empty 96-well plate was placed under each microscope, and acquisition times were measured for different lateral and axial scan steps.

### MCAS image preprocessing

After collecting 3D data from specimens, we registered the images to compensate for lateral shifts observed during axial scanning. Ideally, no lateral movement should occur while performing an axial scan. However, due to minor misalignments between the sample and the MCAS imaging hardware, as well as slight angular differences in the mounted MCAS lenses, lateral shifts were detected. To address this, we measured the lateral shifts for each lens using a custom target containing microspheres of various sizes suspended in a transparent liquid medium, and registered the focal stacks based on this shift information. Finally, we performed a maximum intensity projection for each lateral scan position to reduce dimensionality.

We further investigated how organoid reorientation affects cross-sectional area measurements. Given the spherical morphology of organoids and the potential for positional shifts during media change, it is important to assess whether imaging from different XY positions or rotational angles impacts the accuracy of these measurements. This analysis is shown in Supplementary Figure 1.

### MCAS segmentation and stitching

To segment organoids from the maximum projection images, we utilized Segment Anything for Microscopy (SAM) with the ViT-B encoder, a tool that supports both interactive and automatic segmentation based on the Segment Anything Model [17,18]. After segmentation, any holes within the segmented organoids were filled using contour filling in OpenCV. In experiments involving lateral scanning, image stitching was performed after segmentation using the open-source Hugin codebase [14,33].

### MCAS data analysis

Organoid size was determined by the number of pixels within the segmented regions. Each pixel corresponds to an area of 0.25 µm², calculated as the square of the effective pixel size at the specimen plane. The effective pixel size was obtained by dividing the sensor pixel pitch (1.1 µm) by the objective lens magnification (2.2X). Eccentricity is calculated as the ratio of the distance between an ellipse’s two foci to the length of its major axis. A value of 0 corresponds to a perfect circle, and the closer the value is to 1, the more elongated the ellipse becomes, approaching the shape of a line. Cell confluency was calculated as the total area segmented as cells divided by the area of the well.

Given the large image size (3072 × 3072), traditional dimensionality reduction methods such as PCA are computationally demanding. As an alternative, we utilized CLIP (Contrastive Language–Image Pre-training), a powerful multimodal model that learns semantic relationships and generates numerical embeddings from images. This approach allowed us to reduce dimensionality with CLIP and then apply UMAP to visualize the embeddings in a 2D plot [20,21]. We observed, however, that the background of organoids (e.g., reflected light, intensity variations, debris) influenced the CLIP embedding process. To address this, we implemented mask-aware pooling with ViT-L/14@336, which preserves morphological signals while suppressing background noise. CLIP processes images in patches, and near the end of its pipeline, the patch tokens are aggregated to form an image embedding. Instead of treating all patch tokens equally, we applied mask-aware weighting: for each patch, we computed a weight equal to the fraction of foreground (non-background or segmented organoid) pixels within that patch. The final image embedding was then obtained as a weighted mean of the patch tokens, ensuring that patches dominated by background contributed less to the overall representation.

### iPSC Culture and Maintenance

The iPSC lines were thawed in a 6-well plate coated with hESC-qualified Matrigel (Corning® Matrigel® hESC-Qualified Matrix, LDEV-free, Cat# 354277) in mTeSR basal media (Cat # AJ29845120) supplemented with mTSER supplement (Cat# 1000151549) along with 50nM ROCK inhibitor Chroman-1(Cat# 06673). Cells were fed daily with mTeSR+ media and maintained at 37°C in a 5% CO₂ incubator. The iPSCs were passaged at ∼70% confluency using Versene (Gibco, Cat# 15040-066) and was confirmed to be mycoplasma-free using a PCR detection kit (Cat# 30-1012K).

### Organoid differentiation

Cortical organoids were generated by modifying a reproducible dorsal forebrain differentiation protocol [11]. Neuroectoderm fate was induced via combined Wnt and SMAD inhibition.

On Day 0, human derived iPSCs at ∼70–80% confluency, were dissociated using Accutase (ThermoFisher Scientific, Cat# A1110501), and 9,000 cells per well were seeded into a V-bottom ultra-low attachment 96-well plate (SBIO, MS-9096VZ). Depending on the experiment, cells were either plated in 200 µL of CDM1 media containing 3 µM IWR1, 2 X Chroman, and 5 µM SB431542 for condition 1 or 200 µL of mTeSR+ medium per well (Stemcell technologies, Cat#100-0274) supplemented with 100nM Chroman-1 (MedChem Express, Cat# HY-15392) for conditions 2 and 3. On Day 1, either 50% (condition 2) or 75% (condition 3) of the mTeSR+ media was replaced with CDM1 media (composition: G-MEM; ThermoFisher Scientific #11710-035, KnockOut Serum Replacement; ThermoFisher Scientific #10828-028, MEM Non-Essential Amino Acids Solution; ThermoFisher Scientific #11140-050, Sodium Pyruvate; ThermoFisher Scientific #11360-070, 2-mercaptoethanol; ThermoFisher Scientific, #21985-023,Penicillin/Streptomycin Solution; Gibco #15140-122) containing 3µM Wnt inhibitor, IWR1 (Millipore, #681669), and 5 µM SMAD inhibitor, SB (Stemcell Technologies, #72234) to promote a dorsal telencephalic fate. The organoids were maintained in the 96-well plate until Day 18, with media changes on Days 3, 6, 9, 12, and 15 for all the three conditions. For each media change, 90 µL of the existing CDM1 media in each well was replaced with 100 µL of fresh CDM1 media supplemented with IWR1 and SB. On Day 18, the CDM1 media was completely switched to CDM2 media (composition: DMEM/F-12; ThermoFisher Scientific #11330-032, GlutaMAX; ThermoFisher Scientific #35050-06, N-2; Life Technologies #17502-048, Chemically Defined Lipid Concentration; ThermoFisher Scientific #11905-031, Fungizone; ThermoFisher Scientific #15290-018, Penicillin/Streptomycin Solution; Gibco #15140-122). Further on Day 35 media was switched to CDM3 (composition: DMEM/F-12; ThermoFisher Scientific #11330-032, FBS; VWR #97068085, Chemically Defined Lipid Concentration; ThermoFisher Scientific #11905-031, GlutaMAX; ThermoFisher Scientific #35050-06, N-2; Life Technologies #17502-048, Heparin (Sigma #H3149), Fungizone; ThermoFisher Scientific #15290-018, Penicillin/Streptomycin Solution; Gibco #15140-122) along with 1% matrigel basement membrane matrix (Corning #354234) until D70 when they were transitioned to CDM4 medium (composition: DMEM/F-12; ThermoFisher Scientific #11330-032, FBS; VWR #97068085, Chemically Defined Lipid Concentration; ThermoFisher Scientific #11905-031, GlutaMAX; ThermoFisher Scientific #35050-06, N-2; Life Technologies #17502-048, B-27 supplement; ThermoFisher Scientific #17504-044, Heparin (Sigma #H3149), Fungizone; ThermoFisher Scientific #15290-018, Penicillin/Streptomycin Solution; Gibco #15140-122) with 2% matrigel basement membrane matrix (Corning #354234).

### Seeding media conditions and longitudinal MCAS imaging of cortical organoids

As all iPSC lines did not differentiate well when seeded directly in CDM1, we evaluated the impact of different seeding media on organoid survival. iPSCs were cultured under three experimental conditions as described above. Condition 1: Seed in CDM1 (3 µM IWR1; 2× Chroman-1; 5 µM SB431542). On Days 3/6/9/12/15, replace 90 µL with 100 µL fresh CDM1 + IWR1 + SB431542. Condition 2: Seed in mTeSR+ with 2× Chroman-1; on Day 1 replace 100 µL with CDM1 I + IWR1 + SB431542; then follow Condition 1 exchange schedule. Condition 3: Seed in mTeSR+ with 2× Chroman-1; on Day 1 replace 150 µL with CDM1 + IWR1 + SB431542; then follow Condition 1 exchange schedule. Daily imaging of organoids from Day 1 to Day 17 was performed to assess cross-sectional area and survival, enabling the identification of optimal conditions for organoid culture. In this experiment, lateral scanning was unnecessary, as the organoids were small enough to fit within a single FOV. The total acquisition time was 35 seconds for a 1 × 1 lateral scan with 170 focal stacks acquired at a 10 µm step size, using an exposure time of 50 ms. Eccentricity was calculated as the ratio of the distance between the foci of the ellipse to the length of its major axis. A value of 0 indicates a perfect circle, while a value of 1 corresponds to a line.

### Drug treatment and Imaging protocol for Valproic acid, CHIR99021 and Mitomycin C in cortical organoids

Valproic acid (VPA; Sigma Aldrich, Cat# P4543-10G) was dissolved in a balancing solution containing water and DMSO to achieve final concentrations of 0.6 mM and 3 mM. CHIR99021 (Sigma Aldrich, Cat# SML1046) was dissolved in water and DMSO (Fisher bioreagents Cat# BP231-100, Lot 234276) to achieve final concentrations of 0.625 µM, 1.25 µM and 2.5 µM. Mitomycin C (Sigma Aldrich, Cat# M4287-2MG) was dissolved in water to achieve a final concentration of 0.5 µg/ml, 1 µg/ml and 3 µg/ml. Vehicle controls were matched to the solvent conditions: water/DMSO for CHIR99021 and VPA, and water for Mitomycin C.

The PGP1 iPSC cell line [37] was cultured and maintained following the protocol described in Section 1.2, and cortical organoids were differentiated for 31 days as per Condition 3 detailed in Section iPSC Culture and Maintenance. On Day 18 of differentiation, varying concentrations of VPA (0.6 mM and 3 mM), CHIR (0.625 µM, 1.25 µM, and 2.5 µM) were administered to the organoids in CDM2 medium and media was replaced completely with fresh CDM2 on Day 20. For Mitomycin C, the organoids were treated at varying doses and durations (0.5 µg/mL for 24 hours; 1 µg/mL for both 1 hour and 24 hours and 3 µg/mL for 1 hour) in CDM2 medium. Following the treatment, the media for each condition was completely replaced with fresh CDM2 medium (at the end of the 1-hour or 24-hour treatment) to ensure the complete removal of Mitomycin C. To assess the effects of these treatments, organoids were imaged daily from Day 17 post differentiation (baseline, one day prior to drug administration) to Day 31. For this experiment, a 2 × 2 lateral scan ensured full coverage of the organoids, particularly at later growth stages. The total acquisition time was 132 seconds for a 2 × 2 lateral scan with 170 focal stacks acquired at a 10 µm step size, using an exposure time of 60 ms. We used Welch’s *t*-test (two-tailed) to compare the means between the control and each drug dose, as this test does not assume equal variances between groups.

### Organoid generation with variable GFP positive cell ratios and fluorescent imaging workflow

A constitutively active eGFP expressing lentivirus containing a diverse sequence in the 3’ UTR which can be used for lineage tracing [31] was generated by the UNC Lentiviral vector core using previously described protocols [38]. HEK293T cells were transfected with 8.2 µg of plasmid library (gift from Nowakowski lab at UCSF), 6.8 µg Nodamura B (Addgene #17228), 10 µg packaging plasmid, and 5 µg VSV-G envelope plasmid per 10 cm plate, using the calcium phosphate method. Viral supernatants were collected 24-48 h post-transfection, pooled from 10 plates, and concentrated by ultracentrifugation over a sucrose cushion. Viral titers were quantified by qPCR on target cells, and transduction efficiency was confirmed by GFP expression in organoids.

The PGP1 iPSC cell line was transduced with the STICR virus (1 µL of 1.02 × 10¹⁰ IU) for 24 hours. Post-transduction, the cells were passaged, stabilized, and cryopreserved, hereafter referred to as the STICR PGP1 line. Both the STICR PGP1 and unmodified PGP1 cell lines were thawed, cultured, and maintained following the protocol detailed in Section iPSC Culture and Maintenance. Once the iPSCs reached ∼70% confluency, they were dissociated using accutase and combined in three different ratios: 20%, 40%, and 100%. The 20% condition consisted of 20% STICR PGP1 cells mixed with 80% PGP1 cells, the 40% condition consisted of 40% STICR PGP1 cells mixed with 60% PGP1 cells, and the 100% condition used only STICR PGP1 cells. Cortical organoids were then generated as per condition 3 as described in Section Organoid differentiation and differentiated until Day 7. Organoids were imaged daily, and the percentage of fluorescent cells was quantified for each condition. Because the organoids were sufficiently small to occupy a single FOV, lateral scanning was not required. In addition to brightfield imaging, GFP signals were captured in a separate imaging session. For brightfield imaging, the total acquisition time was 20 seconds for a 1 × 1 lateral scan with 80 focal stacks acquired at a 10 µm step size, using an exposure time of 60 ms. For fluorescence imaging, the total acquisition time was 110 seconds for a 1 × 1 lateral scan with 80 focal stacks acquired at a 10 µm step size, using an exposure time of 200 ms and a digital gain of 4. Fluorescence imaging required a longer exposure time, which prevented the use of continuous vertical stage movement during acquisition due to motion blur, thereby increasing the total acquisition time. As the organoids differentiated, the GFP signal intensified. To avoid image saturation and ensure accurate quantification, different exposure times were used on each day. Segmentation masks of the organoids were used both to calculate organoid size and to quantify fluorescence signal intensity more accurately. This segmentation was essential for background removal, helping to eliminate background signals from the analysis. Signal intensity was calculated by summing the raw pixel values within the segmented organoid regions and normalizing by the exposure time.

### Fluorescence-Activated Cell Sorting (FACS)

Organoids on Day 9 post differentiation were dissociated using the Papain Dissociation System (Worthington Biochemical Corp, Cat# LK003150) with DNase and RNaseOUT (Invitrogen, Cat# 10777-019). Minced organoids (<1 mm pieces) were incubated at 37 °C with papain/DNase for 20-30 min, gently triturated, filtered (40 µm), and centrifuged at 300 × g for 7 min at 4 °C. Viability was assessed by trypan blue, and samples with <70% viability were discarded. Cells (1–10 × 10⁵ per reaction) were fixed using the Evercode Fixation v3 kit (Parse Biosciences, Cat# ECFC3300) according to the manufacturer’s protocol: prefixation, fixation, permeabilization, and stop buffer steps, followed by resuspension in the storage buffer. Samples were frozen at −80 °C in a controlled-rate container until further use.

Single cell suspensions from dissolved organoids were thawed at 37 °C and kept on ice prior to sorting. Cells were stained with DAPI to assess viability. Sorting was performed on a BD FACSAria III (BD Biosciences) using a 100 µm nozzle at 20 psi. Forward scatter, side scatter, and pulse-width gating were applied to exclude debris and doublets (Supplementary Figure 2). GFP positive cells were collected into low-retention tubes containing 500 µL of 1× PBS with 0.01% BSA and RNaseOUT (Invitrogen, Cat# 10777-019). For each sample, we recorded cell viability based on DAPI exclusion, FSC/SSC distributions, doublet exclusion efficiency, the percentage and absolute number of GFP⁺ and GFP⁻ events, and the relative composition of each gated fraction.

### Neuro2a monolayer Cell Culture and MCAS imaging workflow for assessing transfection

Mouse neuroblastoma Neuro2a (N2a) cells (ATCC #CCL-131) were cultured in Dulbecco’s Modified Eagle Medium (DMEM, high glucose, with pyruvate; Thermo Fisher #11995065) supplemented with 10% (v/v) fetal bovine serum (FBS; Thermo Fisher #10437028). Cells were maintained at 37 °C in a humidified atmosphere containing 5% CO2.

For transfection experiments, cells were detached using 0.25% Trypsin-EDTA (Thermo Fisher #25200-056), neutralized with DMEM + 10% FBS, and centrifuged at 300 × g for 5 minutes. The cell pellet was resuspended in fresh DMEM + 10% FBS, and approximately 80,000 cells were seeded per well into a Corning 96-Well, Cell Culture-Treated, Flat-Bottom Microplate (Thermo Fisher #3603). Cells were allowed to adhere overnight prior to transfection. Transfection was carried out using Lipofectamine 3000 (Thermo Fisher #L3000008) following the manufacturer’s instructions. For each well, 0.15 µL of Lipofectamine 3000 was diluted in 5 µL of Opti-MEM (Thermo Fisher #31985070) and vortexed. Separately, 100 ng of plasmid DNA (Addgene plasmid #24526, a gift from Darrell Kotton) was diluted in 5 µL of Opti-MEM containing 0.2 µL of P3000 reagent. The two solutions were mixed in a 1:1 ratio and incubated for 10 minutes at room temperature before being added dropwise to each well. Transfection complexes were prepared as a master mix for all wells to ensure consistency.

This experiment required a 6 × 6 scan to cover the entire field of view of a 96-well plate, with 40 focal stacks acquired at a 5-µm step size. Both brightfield and fluorescence images were captured, with total acquisition times of 345 seconds and 51 minutes, respectively. The exposure times were 60 ms for brightfield and 400 ms for fluorescence. For fluorescence imaging, a digital gain of 6 and an analog gain of 1.06 were applied. The longer exposure time required for fluorescence prevented the use of continuous vertical stage movement during acquisition due to motion blur, thereby increasing the total acquisition time. After collecting the 3D data, the most in-focus plane for each lateral scan position was identified from the brightfield images using a contrast metric based on the Laplacian operator with a 7 × 7 kernel. These in-focus planes were then used for subsequent analysis. For the fluorescence channel, three focal planes were selected prior to performing a maximum intensity projection. The location of the most in-focus plane in each brightfield image stack was used to determine the center plane of the corresponding fluorescence stack. Cell counting was performed on both brightfield and fluorescence images using Cellpose 2.0 [39] with the cyto2 model, and the transfection ratio was calculated by dividing the number of cells detected in the fluorescence images by the number detected in the brightfield images.

### Assessment of Organoid Cross-Sectional Area in 24 well Plate Formats at later differentiation stage

Standard protocols for cortical organoid generation utilize multiple plate formats throughout the differentiation timeline to accommodate increasing organoid size. Initially, organoids are maintained in 96-well plates from Day 1 to Day 18, followed by transfer to 90 mm dishes until Day 35, and subsequently to spinner flasks for long-term culture. As organoids mature, their increasing size poses challenges for imaging in smaller well formats such as 96-well plates. To assess the ability of our imaging system to quantify cross-sectional area in larger, mature organoids, we transferred organoids derived from multiple iPSC lines that had been differentiated to Day 35, 56, 70, or 84 into 24-well plates and imaged them at the corresponding time points. In this experiment, a 7 × 7 scan was used to image a 24-well plate. A total of 40 focal stacks were acquired at a 40 µm step size with an exposure time of 60 ms, and the total acquisition time was 468 seconds. Prior to maximum intensity projection, nine focal planes were selected to improve image contrast. The FOV of the MCAS was slightly smaller than the full area of the 24-well plate, resulting in cropped organoids in the rightmost column. These cropped organoids were excluded from the analysis.

## Declarations

iPSCs were generated from a UNC IRB approved project (IRB: 17-871).

## Data and code availability

Data and code are available at https://doi.org/10.7924/r49k4mh5t.

## Funding

For authors affiliated with Duke University and Ramona Optics Inc.

Research reported in this publication was supported by the National Institute of Mental Health (NIMH) of the National Institutes of Health under Award Number R43MH133521, the Office of Research Infrastructure Programs (ORIP), Office Of The Director, National Institutes Of Health of the National Institutes Of Health and the National Institute Of Environmental Health Sciences (NIEHS) of the National Institutes of Health under Award Number R44OD024879, the National Cancer Institute (NCI) of the National Institutes of Health under Award Numbers R44CA285197 and R44CA250877, a Duke-Coulter Translational Research Grant Award, and a National Science Foundation Award (2036439).

For authors affiliated with University of North Carolina at Chapel Hill

This work was supported by the NIH (R01 MH130441, U01 AG088667, 1R43MH133521-01A1) and the Autism Science foundation Pre-doctoral fellowship (24-005 to R.S).

## Authors’ contributions

K.K., R.S., J.S., and R.H. conceived the idea and initiated the research. K.K., A.C., and K.L. developed the algorithms and performed data analysis. K.K. and A.C. developed customized Gigaviewer. K.K., A.C., J.L., C.D., P.R., A.B., M.H., and R.H. developed the MCAM hardware and acquisition software. With the help of J.P, I.K.,, and J.S., K.K. and R.S. acquired and analyzed the data. R.S. and J.P. generated all the organoids and maintained the well plates in this study. R.S., J.P., performed the single cell fixation and FACS sorting experiments. I.K. prepared and maintained the monolayer experiment plate. K.K., R.S., and F.S. wrote the manuscript and created the figures, with input from all authors. All authors revised the manuscript. J.S., and R.H. supervised the research. All authors have read and agreed to the content.

## Acknowledgements

We are grateful to the UNC Flow Cytometry Core Facility for their assistance with cell sorting using BD FACSAria III. We thank Ryan Delgado and Tomasz J. Nowakowski for generously providing the STICR barcode library used for lentiviral transduction in this study. We also acknowledge the use of Biorender (www.Biorender.com) for figure illustrations and Blender v3.6 (Blender Foundation) for 3D visualizations.

