## Supplementary Figure 1, Supplementary Figure 2 for "Parallelized Brightfield and Fluorescence Imaging of Organoids Using a Scalable Multi-Camera Platform"

### 1. Evaluating the Impact of Organoid Reorientation on Cross-Sectional Area Measurements

The MCAS system acquires images of organoids along the Z-plane. However, given the spherical morphology of organoids and the potential for positional shifts during media changes, it is important to evaluate whether imaging from a different XY perspective or rotational angles affects the measurement of cross-sectional area. To investigate this, we assessed whether the average cross-sectional area of organoids changes following a slight repositioning induced by media disturbance. In this experiment, 187 organoids derived from iPSC were cultured in two 96-well plates and imaged on Day 10 post-differentiation. The total acquisition time per plate was 35 seconds for a 1 × 1 lateral scan with 170 focal stacks acquired at a 10 μm step size, using an exposure time of 60 ms. After the initial imaging, the organoids were gently agitated using a multichannel pipette to stimulate the effect of a media change. Imaging was then repeated to evaluate any differences in cross-sectional area due to organoid reorientation. This approach allowed us to determine the reproducibility and robustness of cross-sectional area measurements across cell lines and imaging angles.

**a**

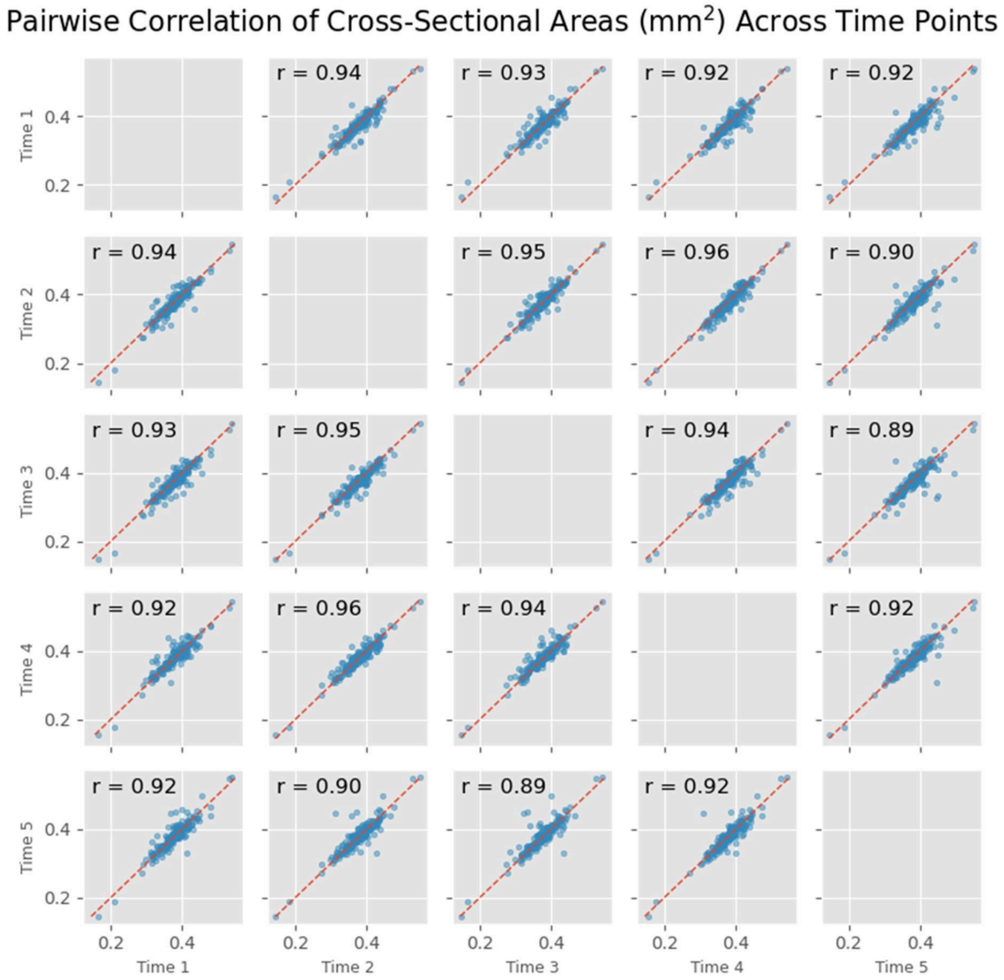


**Supplementary Figure 1. Pairwise correlation of cross-sectional areas (mm$^2$) across time points**.

**a**. Following the initial imaging, the organoids were perturbed with a multichannel pipette to simulate the effects of a media change. Imaging was then performed four additional times to assess whether reorientation influenced cross-sectional area measurements.

### 2. FACS sorting results

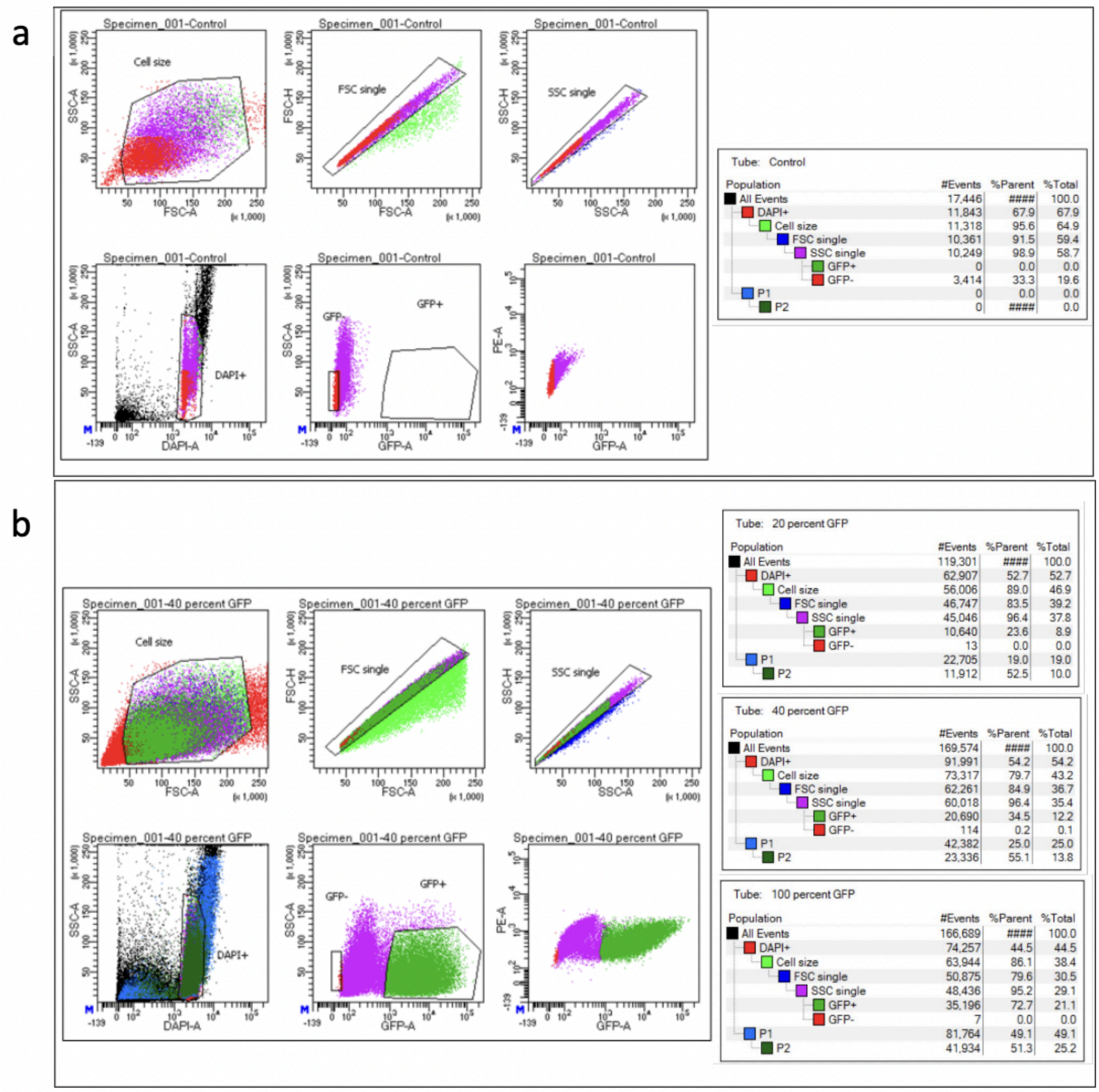


**Supplementary Figure 2. Gating strategy for fluorescence-activated cell sorting (FACS).**

**a**. Representative gating workflow using a negative control sample without GFP expression. Sequential gates were applied to select cells based on size (FSC/SSC), singlet discrimination, and viability (DAPI–), and confirmation of absence of GFP+ events.

**b**. Gating workflow for samples containing GFP+ cells. Cell populations were gated sequentially as in the control, and GFP+ fractions were identified at varying percentages (20%, 40%, and 100% GFP). Plots show the progression of gating from all events to live single cells, with GFP+ and GFP– populations clearly separated. Tables summarize the percentage of events retained at each gating step across control and GFP+ samples.
